# Prefrontal 5-HT_2A_ receptors directly contribute to tic ontogeny: translational evidence

**DOI:** 10.64898/2025.12.29.693270

**Authors:** Easton van Luik, Roberto Cadeddu, Giulia Braccagni, Marco Bortolato

## Abstract

Tourette syndrome (TS) is a neuropsychiatric disorder characterized by motor and vocal tics and frequently accompanied by comorbidities such as obsessive-compulsive disorder (OCD) and pathological aggression. Although approved pharmacotherapies often reduce tic severity, they frequently cause adverse effects and are insufficient for managing comorbid symptoms, underscoring the need for improved treatments. Recent evidence indicates that the serotonin 5-HT_2A_ receptor (5-HT_2A_R) antagonist pimavanserin reduces tic severity and improves quality of life, yet its mechanism of action in TS remains unclear. Here, we combine postmortem human and preclinical approaches to gain insight into the role of 5-HT_2A_Rs in TS. Western-blot analyses of postmortem prefrontal cortex (PFC) samples obtained from individuals with TS revealed a pronounced, male-specific elevation of 5-HT_2A_R protein levels in Brodmann Area (BA) 10. We then tested pimavanserin and the selective 5-HT_2A_R antagonist volinanserin in two mouse models of TS: D1CT-7 transgenic mice and mice with early-life depletion of striatal cholinergic interneurons. Systemic administration of pimavanserin (1–2 mg·kg⁻¹, IP) or volinanserin (0.1–0.3 mg·kg⁻¹, IP) robustly reduced tic-like movements and stereotypies in both models. Local infusion of pimavanserin into the medial PFC, but not the dorsal striatum, recapitulated these effects, indicating a cortical locus of action. Pimavanserin also reduced resident-intruder aggression, but not locomotion or anxiety-like behavior. Together, these findings identify elevated prefrontal 5-HT_2A_Rs as a key mechanistic contributor to TS and a promising therapeutic target for individuals with TS and pathological aggression.

## INTRODUCTION

Tics are repetitive, semivoluntary movements or vocalizations that profoundly impair daily functioning and quality of life. The most disabling tic disorder, Tourette syndrome (TS), is a childhood-onset condition defined by multiple motor and at least one phonic tic persisting for over one year ^1^. TS has a major negative impact on quality of life ^2–4^, due to its detrimental influence on interpersonal relationships, financial stability, and educational attainment ^4,5^. This burden is often compounded by a broad range of comorbid conditions, including obsessive-compulsive disorder (OCD), attention-deficit/hyperactivity disorder (ADHD), anxiety, mood disorders, and disruptive behavior disorders ^6,7^.

Converging evidence from structural and functional neuroimaging studies implicates aberrant cortico-striato-thalamo-cortical (CSTC) circuitry in the pathophysiology of TS ^8,9^. A central feature of this dysfunction is hyperactive dopaminergic signaling within cortico-striatal circuits, as demonstrated by imaging and postmortem studies ^10–14^. Consistent with this mechanism, two of the three medications approved by the US Food and Drug Administration (FDA) for the treatment of TS are first-generation antipsychotics, which reduce tic severity through potent antagonism of dopamine D_2_ receptors (D_2_R) ^15^.

Nevertheless, these compounds impose severe side effects, substantially limiting long-term use ^16–18^, and show limited efficacy in addressing the psychiatric comorbidities of TS. Atypical antipsychotics such as risperidone and olanzapine, which combine D_2_R antagonism with serotonin 5-HT_2A_ receptor (5-HT_2A_R) blockade, are widely used in TS and reduce tics, aggression, and OCD symptoms ^19–24^. However, their broad pharmacology makes it unclear whether their anti-tic efficacy is specifically attributable to 5-HT_2A_R blockade.

A recent open-label pilot study by Billnitzer and Jankovic ^25^ provided the first direct clinical evidence that selective 5-HT2AR antagonism may be effective as a therapy for TS. In this study, twelve adults with TS received pimavanserin, a selective 5-HT_2A_R inverse agonist approved for the treatment of Parkinson’s disease psychosis ^26^. Among the ten completers, pimavanserin produced a statistically significant, albeit modest (12%), reduction in tic severity at week 8 (P = 0.03), along with improvements in clinician- and patient-rated global impressions, quality of life, and obsessive symptoms. Importantly, no serious adverse events were reported. While these findings identify 5-HT_2A_Rs as a compelling therapeutic target, the mechanisms by which 5-HT_2A_Rs influence tic expression and associated behavioral symptoms remain largely unknown. PET studies show globally increased 5-HT_2A_R binding in the cortex of adults with TS ^27^. The prefrontal cortex (PFC) contains among the highest densities of 5-HT_2A_Rs, particularly in layer V pyramidal neurons with major projections to the striatum ^28–30^. Activation of 5-HT_2A_Rs enhances excitatory drive to the striatum, positioning prefrontal receptors as regulators of motor control and inhibition ^29,31,32^. Functional imaging studies show robust PFC engagement during voluntary tic suppression, with greater suppressibility linked to increased prefrontal activity ^33–35^. Together, these findings converge on the PFC as a plausible site of therapeutic action for 5-HT_2A_R antagonists ^25,27^.

To clarify the role of the PFC, we analyzed postmortem tissue from three key brain regions, namely Brodmann Areas (BA) 8, 9, and 10. We also tested the effects of pimavanserin and the highly selective 5-HT_2A_R antagonist volinanserin in two complementary mouse models of TS: (1) D1CT-7 transgenic mice, which exhibit cortical hyperexcitability ^36^, and (2) mice with early-life depletion of striatal cholinergic interneurons (CIN-d) ^37^. We further examined the effects of local intra-PFC versus intrastriatal infusions of pimavanserin and assessed its impact on behavioral paradigms modeling TS comorbidities, including aggression and anxiety.

## MATERIALS AND METHODS

### Human Postmortem Brain Tissue Collection and Donor Information

Frozen PFC tissue samples from Brodmann area (BA) 8, 9, and 10 were obtained from the NIH NeuroBioBank Brain Tissue Resource Center via the Harvard Brain Tissue Repository. Analyses were conducted on postmortem tissue from the BA8 (42.33 ± 12.44 mg), BA9 (30.18 ± 9.55 mg), and BA10 (26.73 ± 3.97 mg) along with corresponding clinical data, including sex, age (38.2 ± 2.8 years), and postmortem interval (PMI, 21.1 ± 1.4 hours). This study involved age- and sex-matched cohorts of thirteen individuals diagnosed with TS (9 males and 4 females), and thirteen control individuals without any psychiatric history (Supplemental Table 1). No differences were observed between groups in terms of age (t(24) = 0.19, P = 0.85) or PMI (t(24) = 0.03, P = 0.97). All samples were collected in accordance with institutional protocols of the participating Brain Tissue Repositories. Each brain was coronally sectioned, rapidly frozen, and stored at −80°C. The superior and middle frontal gyri (BA8), dorsolateral PFC (BA9), and anterior PFC (BA10) were identified using established cortical landmarks, and white matter was excluded from dissection. No macroscopic atrophy was observed in any sample. Informed consent for brain donation and research use was obtained from next of kin in compliance with all legal and ethical requirements. TS diagnoses were confirmed by experienced research clinicians through structured interviews with family members and comprehensive review of medical records, while control cases were verified to be free of psychiatric or neurological disorders.

### Immunoblotting Procedures

Tissues were stored at −80°C until assayed. Approximately 30–40 mg of tissue per sample was homogenized in RIPA buffer (20 mM Tris-HCl pH 7.5, 150 mM NaCl, 1 mM EDTA, 1 mM EGTA, 1% NP-40, 1% sodium deoxycholate, 2.5 mM sodium pyrophosphate, 1 mM β-glycerophosphate, 1 mM Na_3_VO_4_, 1 μg·ml⁻¹ leupeptin, and protease inhibitor cocktail). After sonication (QSONICA Sonicator, Newtown, CT, USA) for 10 s on ice, homogenates were centrifuged at 20,800 *g* (Eppendorf 5147R) at 4°C. Protein concentrations were determined using the DC protein assay (Bio-Rad Laboratories, Hercules, CA).

Equal protein loads (50–100 μg) were run in triplicate on 4–15% Criterion™ TGX Stain-Free™ precast gels (Bio-Rad) and transferred to nitrocellulose membranes. Membranes were visualized under UV light to quantify total-lane protein as a loading control. After blocking in 3% BSA/TBS-T, membranes were incubated overnight at 4°C with rabbit polyclonal anti-5-HT_2A_R antibody (#ab66049, 1:500 dilution, Abcam, Cambridge, UK). After TBS-T washes, membranes were incubated with goat anti-rabbit HRP-conjugated secondary antibody (#31462, 1:10,000 dilution, ThermoFisher, Waltham, MA, USA) for 90 min at room temperature. Signal detection was performed using Clarity Western ECL substrate (Bio-Rad) and imaged via ChemiDoc™ XRS+ (Bio-Rad). Bands were quantified in arbitrary units using Image Lab software (Bio-Rad) and normalized to total-lane protein.

To verify cytoarchitectural integrity, membranes were stripped and reprobed with anti–β-actin antibody (#sc47778, 1:1000, Santa Cruz Biotechnology, Dallas, TX, USA). No differences in β-actin levels were detected between diagnostic groups, validating normalization procedures.

### Animals

D1CT-7 transgenic mice (C.Cg-Tg(DRD1-ctxA)7Burt/J, strain ID: #008367, Jackson Laboratory, Bar Harbor, ME, USA) and wild-type (WT) littermates were maintained on a 50% C57BL/6 and 50% Balb/c background and genotyped as previously described ^38^. Striatal cholinergic interneuron depleted (CIN-d) mice were generated from offspring of homozygous ChAT-Cre males (B6;129S6-ChAT^tm2(cre)Lowl/J^, strain ID: #006410, Jackson Laboratory) crossed with C57BL/6J females (Jackson Laboratory #000664), following previously described procedures ^37^. Briefly, at postnatal day (PND) 4, heterozygous pups were cryoanesthetized and received bilateral intracranial microinjections of an adeno-associated virus serotype 5 (AAV5) vector expressing the simian diphtheria toxin receptor under a lox-stop-lox cassette or an inactive control construct (coordinates: AP −0.5 mm; ML ±1.6 mm; DV −2.8 mm). At PND 18, animals received a single intraperitoneal injection of diphtheria toxin (1 μg·kg⁻¹) to induce selective partial ablation of Cre-recombinase–expressing cholinergic interneurons (CINs).

Mice were weaned at PND 21 and group-housed (3 per cage) under standard housing conditions (22°C, 12-h light/dark cycle; lights on 6:00 AM–6:00 PM) with *ad libitum* access to food and water. All experiments were performed in adult mice (60-90 day-old) between 9:00 AM and 4:00 PM. Animal numbers were determined by power analysis based on prior effect sizes. Procedures complied with NIH guidelines and were approved by the Institutional Animal Care and Use Committees at University of Utah and University of Florida. Only male mice were used, based on prior evidence of male-specific behavioral alterations and the strong male predominance of TS ^39,40^.

### Surgical Procedures

For targeted drug delivery, adult mice were anesthetized with xylazine/ketamine (20/80 mg·kg⁻¹, IP) and implanted with bilateral guide cannulae (26 gauge; P1 Technologies Inc., Roanoke, VA, USA) targeting either the medial PFC (AP: + 1.94 mm, ML: ± 0.35 mm, DV: −1.30 mm) and the caudate-putamen (CPu; AP: + 1.2 mm, ML: ± 1.6 mm, DV: 3.8 mm).

### Drugs

Pimavanserin and volinanserin (Sigma-Aldrich, St. Louis, MO) were dissolved in a 2.5% DMSO, 2.5% Tween-80, and 0.9% NaCl for IP administration (10 ml·kg⁻¹). Both volinanserin (0.1–0.3 mg·kg⁻¹, IP) and pimavanserin (1–2 mg·kg⁻¹, IP) were administered 30 min before behavioral testing. For intracerebral microinfusions, pimavanserin (1µg in 0.5µl of DMSO) was infused directly into the medial PFC or CPu 10 min before testing. 2,5-Dimethoxy-4-iodoamphetamine (DOI) hydrochloride (Sigma-Aldrich, St. Louis, MO) was dissolved in 0.9% NaCl and injected IP (10 ml·kg⁻¹) 30 min before behavioral testing.

### Behavioral Procedures

Male mice were tested using a comprehensive behavioral battery to assess tic-like, anxiety-related, and aggressive phenotypes relevant to TS and its comorbidities. The behavioral testing battery included:

#### Spatial Confinement and assessment of tic-like behaviors

Tic-like jerks and stereotypic grooming were measured under spatial confinement, a mild stressor that elicits robust increases in these responses in both models ^37,41^. Mice were placed in a clear, bottomless Plexiglas cylinder (10 cm × 30 cm) positioned in their home cage for 40 min. The final 30 min were video-recorded for analysis. Behaviors were scored by observers blind to experimental conditions. Each mouse was tested only once to avoid carry-over stress effects.

#### Open Field Test

Locomotor activity was tested as previously described ^42^. Testing was conducted in a black Plexiglas open-field arena (40 × 40 × 40 cm). Mice were placed in the center of the arena, and spontaneous locomotor activity was recorded for 10 min. Following each session, the arena was cleaned with 70% ethanol to remove odor cues from previous subjects and allowed to dry for 10 min. Behavioral recordings were analyzed using automated tracking software (EthoVision; Noldus Instruments, Wageningen, The Netherlands). The arena floor was divided into two zones of equal area (800 cm²): a central square and a peripheral zone. Outcome measures included total distance traveled (index of locomotor activity) and the percentage of total time spent in the central zone (index of anxiety-like behavior) ^43,44^.

#### Elevated Plus Maze

Anxiety-like behavior was assessed using an elevated plus-maze ^45^ with two open (35 × 6 cm) and two closed (35 × 6 × 21 cm) arms elevated 74 cm from the floor. Mice were placed in the central platform, and time spent in open arms (four paws criterion) was expressed as a percentage of total time.

#### Resident–Intruder Test

Aggression was evaluated using the resident–intruder paradigm^46^. Resident males were pair-housed with females for 14 days prior to testing to promote territoriality. One hour before testing females were removed. Following a 20-min confinement period, an unfamiliar intruder (matched for age, sex, and weight ± 10%) was introduced for 5 min. Attack frequency, latency, and duration were scored by blinded observers.

### Statistical Analyses

Human postmortem data were analyzed using two-way analyses of variance (ANOVA), where diagnosis and sex were included as between-subject factors. Two-way ANOVA were additionally used for animal experiments, with model and treatment serving as factors. Where significant main effects or interactions were detected, Tukey’s post hoc tests were conducted. All data are expressed as mean ± SEM, and statistical significance was set at P < 0.05.

## RESULTS

### 5-HT_2A_ Receptor Expression Is Elevated in the BA10 of Male TS Patients

Western-blot analyses were conducted on postmortem PFC tissue encompassing BA8, BA9, and BA10 from individuals with TS and unaffected controls to quantify 5-HT_2A_ receptor expression. Two-way ANOVAs were performed with diagnosis (TS vs. controls) and sex (male vs. female) as between-subject factors. Analyses of BA8 showed higher 5-HT_2A_R levels in males irrespective of diagnostic status (Fig. 1A), with no significant differences between individuals with TS and controls. In BA9, 5-HT_2A_R expression did not differ across sex or diagnosis (Fig. 1B). In contrast, BA10 samples exhibited a marked increase in 5-HT_2A_R levels in males with TS compared with both control subjects and females with TS (Fig. 1C). However, no differences were observed among female subjects (Fig. 1C). Taken together, these findings indicate that TS is associated with a male-specific elevation of 5-HT_2A_R expression selectively in BA10, but not in other PFC regions.

**Fig. 1.**
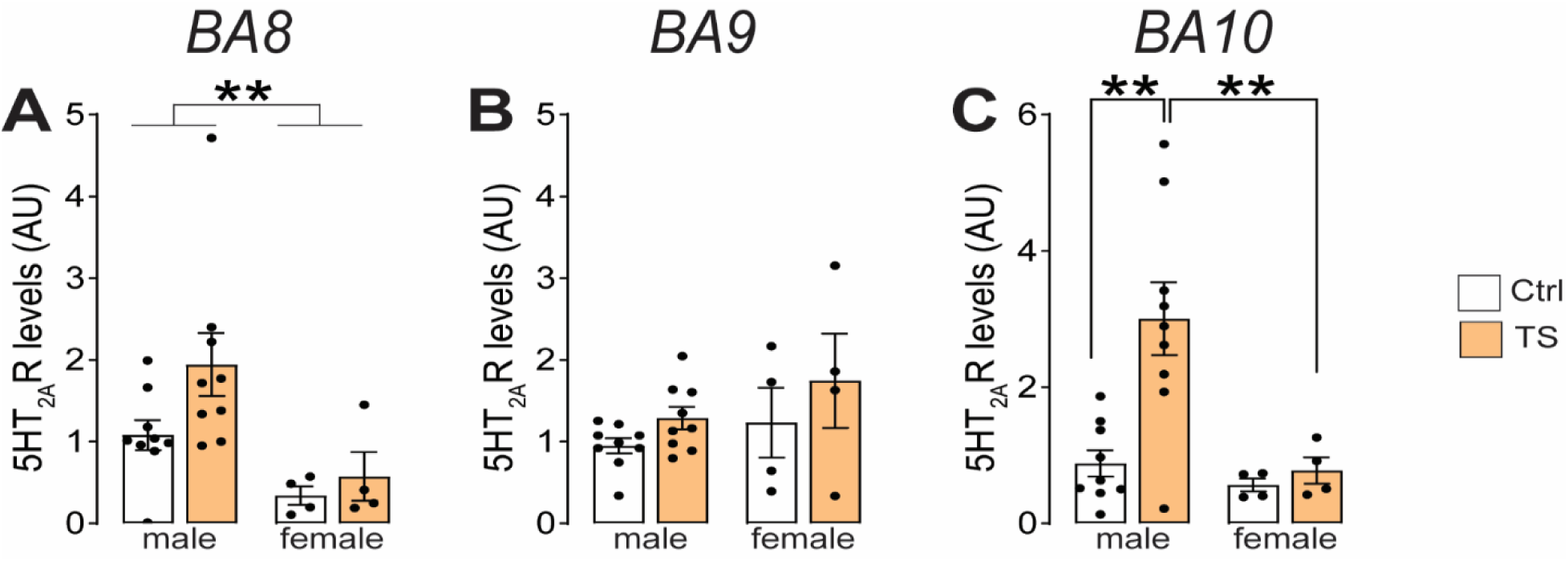
5-HT_2A_R protein expression is selectively elevated in Brodmann Area (BA) 10 of male patients with Tourette syndrome (TS). Western-blot analysis of 5-HT_2A_R protein levels in postmortem prefrontal cortex tissue from TS patients and unaffected controls (Ctrl). (A) Two-way ANOVA of 5-HT_2A_R expression in BA8 revealed a significant effect of sex (F(1,22) = 9.56, P = 0.005), but no significant effects of diagnosis (F(1,22) = 2.58, P = 0.12) or interaction (F(1,22) = 0.85, P = 0.37). (B) Two-way ANOVA of 5-HT_2A_R expression in BA9 showed no significant main effects or interaction (sex: F(1,22) = 2.06, P = 0.17; diagnosis: F(1,22) = 2.71, P = 0.11; interaction: F(1,22)=0.11, P = 0.74) (C) In BA10, analysis revealed significant main effects of diagnosis (F(1,22) = 7.00, P = 0.01) and sex (F(1,22) = 8.31, P = 0.009), as well as a diagnosis x sex interaction (F(1,22) = 4.66, P = 0.042). Post-hoc analysis with Tukey’s correction indicated that male TS patients exhibited significantly higher 5-HT_2A_R receptor expression compared to male control subjects and female TS subjects. Data were analyzed by two-way ANOVA with diagnosis and sex as factors. Symbols: **, P < 0.01 for the comparisons indicated by brackets. Individual points represent independent samples and all data are shown as mean ± SEM.

### Systemic 5-HT_2A_R Antagonism Suppresses Tic-like Behaviors in two different TS mouse models

To determine whether blockade of 5-HT_2A_Rs ameliorates tic-like behaviors, we administered the selective 5-HT_2A_R antagonists volinanserin (0.1–0.3 mg·kg⁻¹, IP) and pimavanserin (1–2 mg·kg⁻¹, IP) in two validated mouse models of TS: the D1CT-7 transgenic line, characterized by cortical hyperexcitability ^47^, and the CIN-d model, which exhibits early-life depletion of striatal cholinergic interneurons ^37^. Both models were tested under spatial confinement, a mild stressor that reliably elicits tic-like jerks and/or excessive grooming stereotypies across these and other animal models of TS ^37,38,41^. D1CT-7 mice displayed a higher frequency of tic-like jerks compared with WT littermates, and this phenotype was normalized by volinanserin treatment (Fig. 2A). In contrast, grooming behavior did not differ significantly between D1CT-7 and WT mice and was not significantly affected by volinanserin in this model (Fig. 2B). In contrast to our previous findings ^37^, no significant difference in baseline jerk frequency was detected between CIN-d mice and controls. Nevertheless, the lower dose of volinanserin (0.1 mg·kg⁻¹) significantly reduced jerk frequency irrespective of CIN status (Fig. 2C). As expected, CIN-d mice exhibited a marked increase in grooming behavior, which was attenuated by both doses of volinanserin (Fig. 2D).

**Fig. 2.**
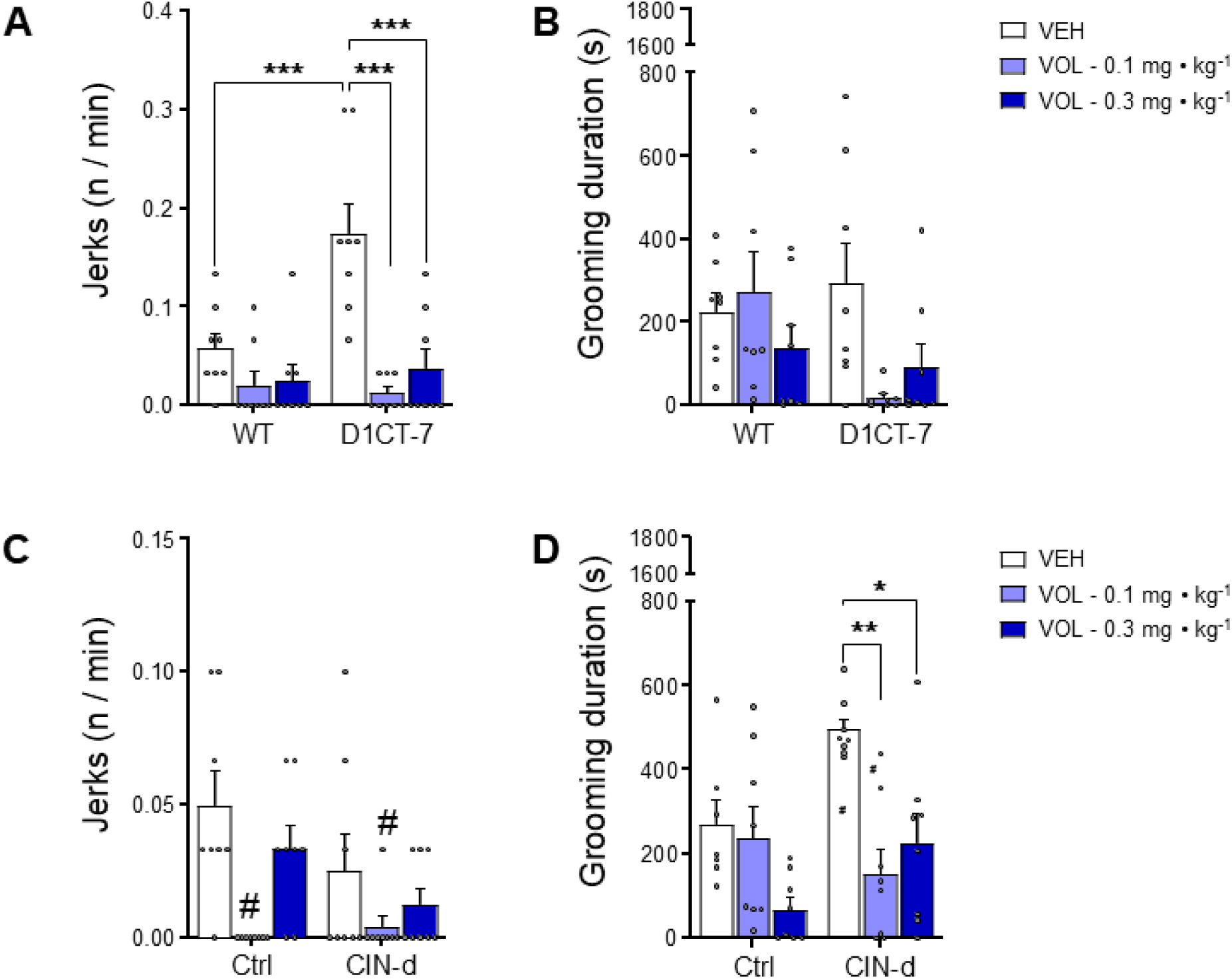
Systemic administration of volinanserin (VOL; 0.1–0.3 mg·kg⁻¹, IP) suppresses tic-like behaviors. Effects of VOL or vehicle (VEH) on tic-like jerk frequency (A,C) and grooming duration (B,D) in D1CT-7 (A-B) and CIN-d mice (C-D). (A) In D1CT-7 mice, analysis of jerks revealed main effects of treatment (F(2, 42) = 17.45, P < 0.0001) and genotype (F(1, 42) = 7.28, P = 0.01), and a significant genotype × treatment interaction (F(2, 42) = 6.71, P = 0.003). (B) VOL treatment did not alter grooming duration in D1CT-7 mice (genotype: F (1,42) = 2.02, P = 0.16; treatment: F (2,42) =2.66, P = 0.08; interaction F (2,42) =3.10, P = 0.06). (C) In CIN-d mice, analysis of jerks revealed a main effect of treatment (F(2, 42) = 7.90, P = 0.0012). No other significant effects were detected (CIN status: F (1,42) =3.61, P = 0.06; interaction: F (2.42) =1.55, P = 0.22). (D) VOL treatment reduced grooming in CIN-d mice (CIN status: (F(1, 41) = 4.81, P = 0.03; treatment (F(2, 41) = 9.71, P = 0.0004); interaction (F (2, 41) = 4.25, P = 0.02). All experiments were performed in male animals with model and treatment as factors. Data were analyzed by two-way ANOVA followed by post-hoc analyses with Tukey’s correction. All data are shown as means ± SEM. *, P < 0.05; **, P < 0.01; ***, P < 0.001. #, P < 0.001 for the post-hoc comparisons of VOL 0.1 mg·kg⁻¹ vs VEH and VOL 0.1 mg·kg⁻¹ vs VOL 0.3 mg·kg⁻¹ (calculated on main effect).

We next assessed the effects of pimavanserin in both mouse models. In D1CT-7 mice, tic-like jerk frequency was higher than in vehicle-treated WT controls, and both doses of pimavanserin significantly reduced jerk frequency in D1CT-7 mice, with no significant effects observed in treated WT animals (Fig. 3A). In line with the effects observed with volinanserin, no significant differences in grooming behavior or treatment-related effects were detected in D1CT-7 mice (Fig. 3B).

**Fig. 3.**
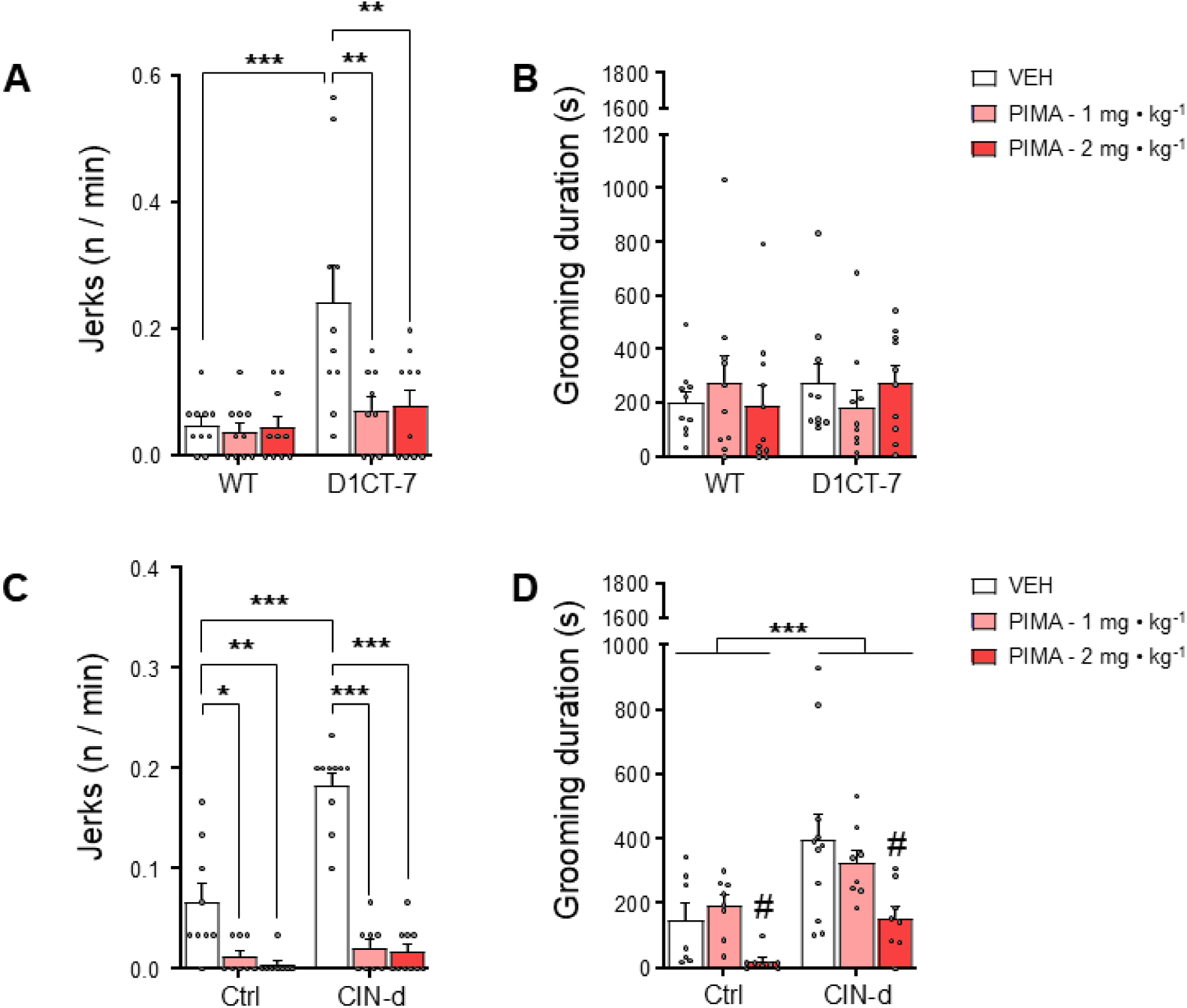
Systemic administration of pimavanserin (PIMA; 1–2 mg·kg⁻¹, IP) suppresses tic-like behaviors. Effects of PIMA or vehicle (VEH) on tic-like jerk frequency (A,C) and grooming duration (B, D) in D1CT-7 (A-B) and CIN-d (C-D) mice. (A) In D1CT-7 mice, analysis of jerks revealed main effects of genotype (F(1, 53) = 13.06, P = 0.0007), treatment (F(2, 53) = 5.81, P = 0.005), and a significant genotype × treatment interaction (F(2, 53) = 4.93, P = 0.01). (B) PIMA treatment did not alter grooming duration in D1CT-7 mice (genotype: F(1,55)= 0.12, P = 0.73; treatment: F(2,55)=0.005, P = 0.99; interaction F(2,55) =1.03, P = 0.36) (C) In CIN-d mice, analysis of jerks revealed main effects of CIN-status (F(1, 47) = 25.56, P < 0.0001), treatment (F(2, 47) = 70.25, P < 0.0001), and a significant interaction (F(2, 47) = 15.86, P < 0.0001). (D) Analyses of grooming revealed main effects of CIN status (F(1, 44) = 14.95, P = 0.0004) and PIMA treatment (F(2, 44) = 7.07, P = 0.002), with no significant interaction (F(2, 44) = 0.79, P = 0.46). All experiments were performed in male animals with model and treatment as factors. Data were analyzed by two-way ANOVA followed by post-hoc analyses with Tukey’s correction. All data are shown as means ± SEM. *, P < 0.05; **, P < 0.01; *** P < 0.001 for the comparison indicated by brackets. #, P < 0.01 for the post-hoc comparisons of PIMA 2 mg·kg⁻¹ vs PIMA 1 mg·kg⁻¹ and PIMA 2 mg·kg⁻¹ vs VEH (calculated on main effect).

In CIN-d mice, pimavanserin significantly reduced tic-like jerk frequency. Vehicle-treated CIN-d mice showed higher jerk frequency than vehicle-treated controls, and pimavanserin reduced these movements at both tested doses (Fig. 3C). Analysis of grooming duration revealed significant main effects of CIN status and treatment (Fig. 3D), indicating increased grooming in CIN-d mice relative to controls and a reduction in grooming duration following pimavanserin treatment irrespective of CIN status.

Taken together, these results indicate that selective 5-HT_2A_R antagonism effectively suppresses tic-like behaviors across mechanistically distinct mouse models of TS.

### 5-HT_2A_R Activation Induces Tic-like Behaviors in CIN-d Mice

To determine whether 5-HT_2A_R activation is sufficient to elicit tic-like behaviors, we administered low doses of the selective agonist 2,5-dimethoxy-4-iodoamphetamine (DOI; 0.05 mg·kg⁻¹, IP) to WT and CIN-d mice. In the absence of the spatial confinement stressor, DOI selectively increased jerk frequency in CIN-d mice, whereas no changes were observed in WT mice (Fig. 4A). Conversely, no effect was found on grooming behavior (Fig. 4B). Notably, because this assessment was conducted without confinement-induced stress, these results indicate that increased 5-HT_2A_R signaling is sufficient to precipitate tic-like jerks, but not stereotypies, supporting the idea that this receptor may be sufficient for brief, sudden movements of the head or the trunk, but not for more complex behavioral sequences.

**Fig. 4.**
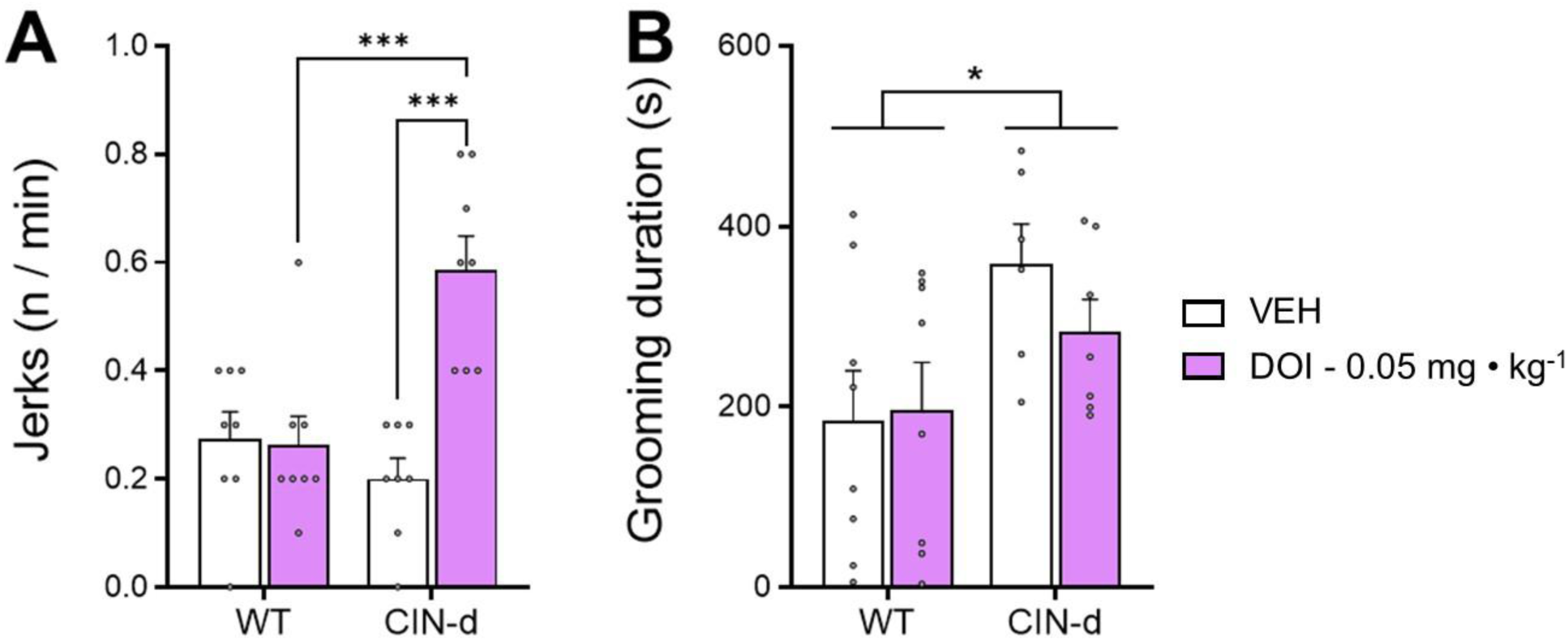
Subthreshold dose of DOI (0.05 mg·kg⁻¹, IP) selectively induces tic-like jerks in CIN-d mice. Effect of the 5-HT_2A_R agonist DOI on tic-like jerk frequency in WT and CIN-d mice. (A) Analysis of jerk frequency following DOI treatment identified significant effects of CIN status (F(1, 25) = 10.00, P = 0.004), treatment (F(1, 25) = 11.58, P = 0.002), and interaction (F(1, 25) = 13.29, P = 0.001). (B) Grooming analysis showed a significant main effect for CIN status (F(1, 25) = 6.89, P = 0.015), with no significant effects of treatment or interaction (treatment: F(1, 25) = 0.39, P = 0.54); interaction: F(1, 25) = 0.74, P = 0.40). All experiments were performed in male animals with model and treatment as factors. Data were analyzed by two-way ANOVA followed by post-hoc analyses with Tukey’s correction. All data are shown as mean ± SEM. *, P < 0.05 and *** P < 0.001 for the comparisons indicated by brackets.

### Region-Specific Cortical Pimavanserin Infusion Reduces TS-Related Motor Abnormalities

To identify the neuroanatomical locus underlying the anti-tic effects of 5-HT2AR blockade, pimavanserin (1 µg per side) was locally infused into either the medial PFC or the CPu of D1CT-7 and CIN-d mice. Infusion of pimavanserin into the medial PFC significantly reduced jerk frequency and grooming duration in both models; however, these effects were not model-specific, as comparable reductions in tic-related behaviors were also observed in control animals (Fig. 5A-D). Notably, CIN-d mice did not show an elevation in either tic-like jerks or grooming stereotypies, likely reflecting the impact of intracranial surgery, which appeared to increase baseline responses in WT controls. In contrast, intra-CPu infusion of pimavanserin failed to alter either jerk frequency or grooming duration in D1CT-7 or CIN-d mice (Fig. 5E-H). Collectively, these findings identify the medial PFC as the critical locus mediating the behavioral effects of pimavanserin, consistent with the region-specific upregulation of cortical 5-HT_2A_R observed in postmortem tissue from individuals with TS.

**Fig. 5.**
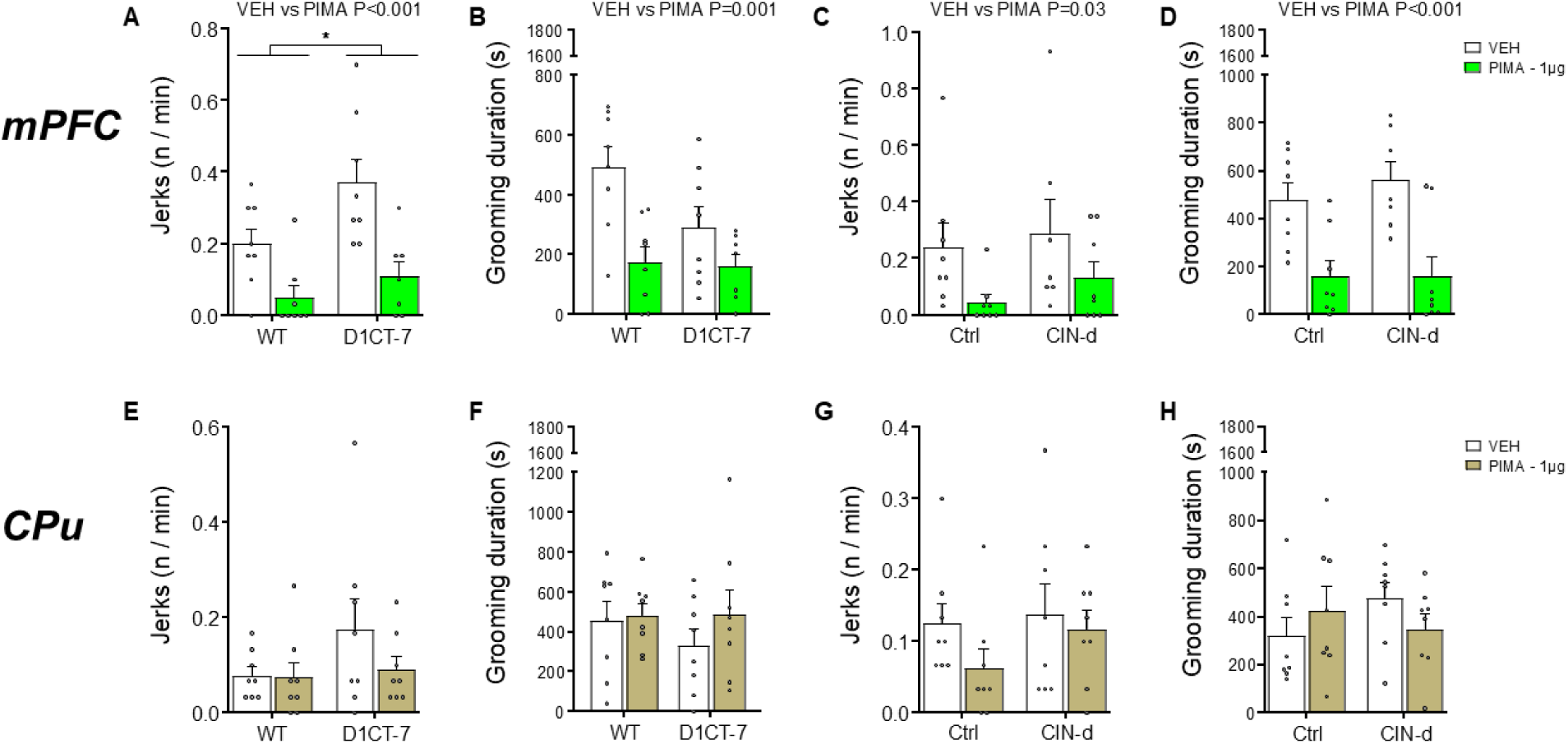
Region-Specific Cortical pimavanserin Infusion Reduces TS-Related Motor Abnormalities. Effects of pimavanserin (PIMA, 1 µg/side) or vehicle (VEH) infused into the medial prefrontal cortex (mPFC, A-D) or caudate-putamen (CPu, E-H) on tic-like jerk frequency and grooming duration in D1CT-7 and CIN-d mice. (A) Analysis of D1CT-7 mice following intra-mPFC PIMA infusion revealed significant effects on jerk frequency for treatment (F(1,27) = 18.89, P = 0.0002) and genotype (F(1,27) = 5.92, P = 0.02), while no interaction was detected (F(1,27) = 1.38, P = 0.25). (B) Grooming duration in D1CT-7 was significantly affected by treatment alone (F(1,27) = 13.58, P = 0.001) irrespective of genotype (genotype: F(1,27) = 3.21, P = 0.08; interaction: F(1,27) = 2.38, P = 0.13). (C) In CIN-d mice following intra-mPFC PIMA infusion, analysis of both jerk frequency and grooming revealed a significant main effect of treatment irrespective of CIN status (jerk frequency: treatment, F(1,27) = 5.37, P = 0.03, CIN status, F(1,27) = 0.80, P = 0.38, interaction, F(1,27) = 0.06, P = 0.80; grooming duration: treatment, F(1,27) = 23.83, P < 0.0001, CIN status, F(1,27) = 0.30, P = 0.59, interaction, F(1,27) = 0.31, P = 0.58). (E–H) No significant effects were detected for CPu infusions irrespective of TS-mouse model (D1CT-7 jerk frequency: genotype, F(1,28) = 2.03 and P = 0.16, treatment, F(1,28) = 1.23 and P = 0.28, interaction, F(1,28) = 1.01 and P = 0.32; D1CT-7 grooming duration: genotype, F(1,28) = 0.38 and P = 0.54, treatment, F(1,28) = 1.00 and P = 0.33, interaction, F(1,28) = 0.49 and P = 0.49; CIN-d jerk frequency: CIN status, F(1,28) = 1.08 and P = 0.31, treatment, F(1,28) = 1.71 and P = 0.20, interaction, F(1,28) = 0.43 and P = 0.52; CIN-d grooming duration: CIN status, F(1,28) = 0.28 and P = 0.60, treatment, F(1,28) = 0.02 and P = 0.88, interaction, F(1,28) = 2.37 and P = 0.13). All experiments were performed in male animals with model and treatment as factors. Data were analyzed by two-way ANOVA followed by post-hoc analyses with Tukey’s correction. All data are shown as means ± SEM. *, P <0.05 for the comparison indicated by bracket.

### Pimavanserin reduces aggressive behaviors, but not locomotion or anxiety-like responses

Given the established involvement of 5-HT_2A_Rs in aggression, we examined whether pimavanserin modulates behavioral phenotypes relevant to these domains. To this end, the D1CT-7 mouse model was selected, as it features spontaneous increase in aggression ^47^.

Aggressive behavior was quantified using the resident–intruder paradigm. We confirmed that D1CT-7 mice exhibited more aggressive behavior relative to WT littermates. Pimavanserin (2 mg·kg⁻¹, IP) significantly reduced both the number (Fig. 6A) and duration (Fig. 6B) of attack bouts. Collectively, these data indicate that 5-HT_2A_R antagonism attenuates the aggressive phenotype of D1CT-7 mice. As shown in Fig. 6C, WT and D1CT-7 mice exhibited similar levels of locomotor activity, and pimavanserin did not influence these effects. Interestingly, D1CT-7 mice showed less thigmotaxis than their WT counterparts (Fig. 6D), but this difference was not affected by pimavanserin treatment, confirming that the antiaggressive effects of this drug were not due to locomotor changes. Similar to these data, D1CT-7 mice showed reduced baseline anxiety-like behavior in the elevated plus maze, but no significant effect of pimavanserin was found (Supplemental Fig. 1). Collectively, these data indicate that the effects of pimavanserin on TS-related phenotype in these animal models are not secondary to nonspecific motor impairment or anxiolysis.

**Fig. 6.**
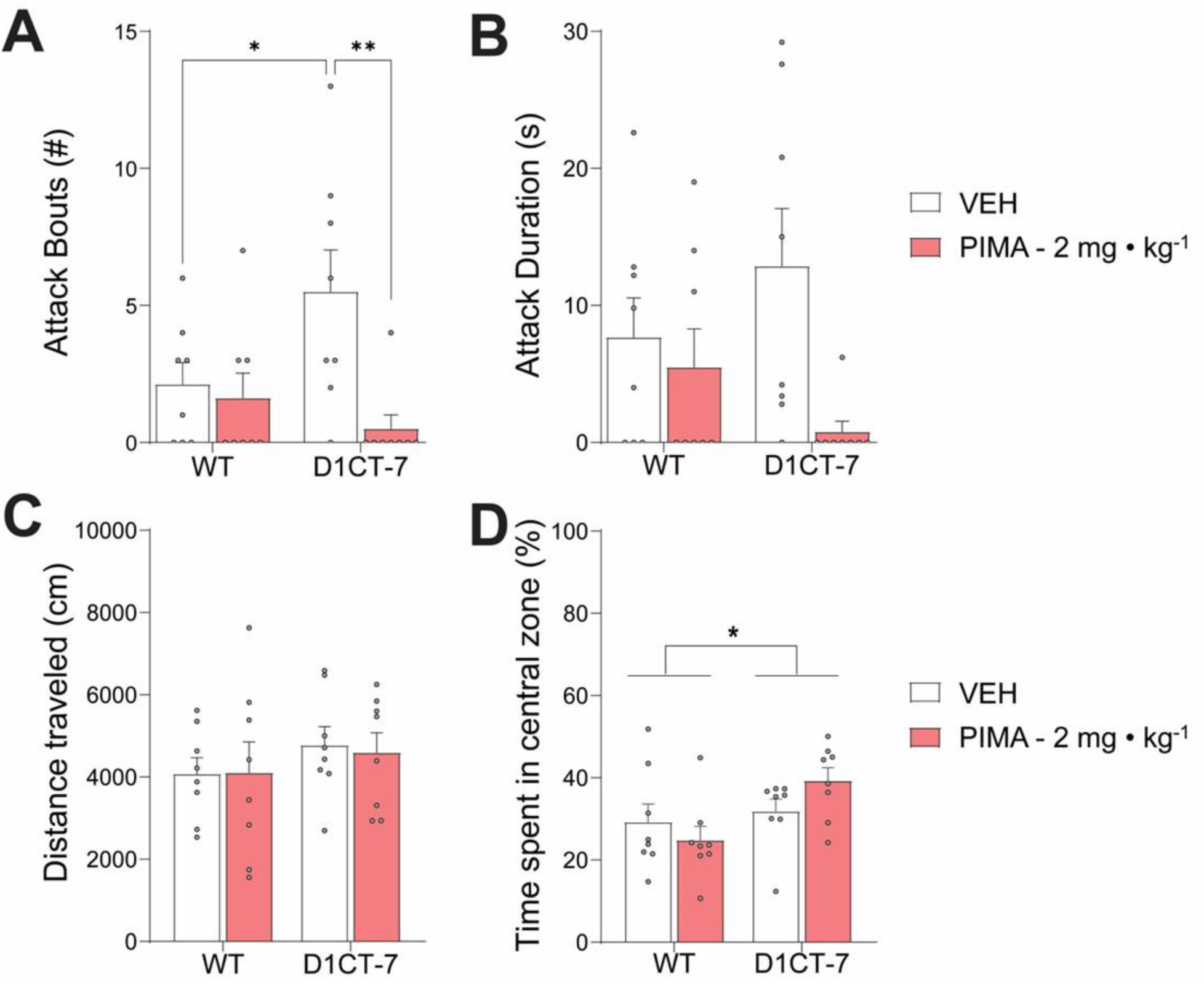
Pimavanserin (PIMA; 2 mg·kg⁻¹) reduces aggression without impairing locomotor activity. Effects of PIMA or vehicle (VEH) on resident-intruder aggression (A-B) and locomotory activity in the Open Field test (C-D). (A) Analysis of the number of attacks showed a significant treatment effect (F(1, 28) = 7.54, P = 0.01) and genotype × treatment interaction (F(1, 28) = 5.05, P = 0.03), but no significant effect for genotype (F(1,28) = 1.26, P = 0.27). (B) Attack duration analysis showed a significant main effect of treatment (F(1, 28) = 5.95, P = 0.02), but not genotype (F(1, 28) = 0.001, P = 0.94) nor interaction (F(1, 28) = 2.88, P = 0.10). (C-D) In the Open Field test, total distance traveled in D1CT-7 mice showed no significant effects of genotype (F(1, 28) = 1.22, P = 0.28), treatment (F(1, 28) = 0.02, P = 0.89), or interaction (F(1, 28) = 0.04, P = 0.85). However, analysis of time spent in the central zone revealed a significant main effect of genotype (F(1, 28) = 5.85, P = 0.02) but no main effect of treatment (F(1, 28) = 0.18, P = 0.67) or interaction (F(1, 28) = 2.80, P = 0.11). All experiments were performed in male animals with model and treatment as factors. Data were analyzed by two-way ANOVA followed by post-hoc analyses with Tukey’s correction. All data are shown as means ± SEM. *, P < 0.05, **, P<0.01 for comparisons indicated by brackets.

## DISCUSSION

The present study yielded four main findings that converge on a pivotal role for 5-HT_2A_Rs in the anterior PFC in TS. First, postmortem analyses revealed a pronounced and regionally restricted increase in 5-HT_2A_R protein levels in the anterior PFC of male individuals with TS. Second, consistent with recent clinical findings ^25^, pimavanserin (as well as the highly selective 5-HT_2A_R antagonist volinanserin) robustly suppressed tic-like motor behaviors and stereotypies in both D1CT-7 and CIN-d mice, two distinct models capturing complementary mechanisms of TS pathophysiology. Conversely, 5-HT_2A_R activation with low doses of a potent agonist produced a significant increase in tic-like jerks, but not grooming stereotypies, in CIN-d mice, supporting the interpretation that 5-HT_2A_R signaling is both necessary and sufficient for the expression of tic-like jerks, but only necessary for more complex stereotyped behaviors. Third, the effects of systemic pimavanserin were reproduced by direct infusions of this drug into the medial PFC, but not the CPu, pointing to the medial PFC as the critical locus of therapeutic action. Fourth, pimavanserin reduced resident-intruder aggression in D1CT-7 mice without altering locomotor activity or anxiety-related measures, supporting the potential of 5-HT_2A_R antagonists to ameliorate clinically burdensome comorbid domains often refractory to current treatments. Collectively, these findings provide convergent translational evidence that excessive or dysregulated 5-HT_2A_R signaling within the PFC contributes to tic ontogeny and represents a promising therapeutic target for TS, particularly in individuals with comorbid disruptive disorders.

A mechanistic framework for these effects is supported by the cellular localization and signaling properties of 5-HT_2A_Rs within the PFC. 5-HT_2A_Rs are densely expressed in layer V pyramidal neurons projecting to the midbrain and striatum ^29,48^. Activation of these receptors depolarizes pyramidal cells and amplifies glutamatergic transmission via canonical Gq-coupled pathways engaging phospholipase C signaling and intracellular calcium mobilization ^31,49,50^. Electrophysiological studies confirm that selective 5-HT_2A_R activation increases the frequency and amplitude of asynchronous spontaneous excitatory postsynaptic currents (sEPSCs) in these neurons ^31,49^. Because tics can be conceptualized as failures of inhibitory gating that permit release of context-inappropriate motor fragments, excessive 5-HT_2A_R-driven excitatory gain in PFC output neurons may destabilize corticofugal excitation-inhibition balance. The stochastic, burst-like nature of these excitatory events also offers a mechanistic substrate for the waxing-waning and semi-voluntary quality of tics, which can be transiently suppressed yet recur when inhibitory resources are depleted or during heightened arousal. Cortical 5-HT_2A_R antagonism may therefore attenuate tic expression by lowering excitatory gain impinging on downstream CSTC nodes. This cortical mechanism also offers a coherent bridge between serotonergic and dopaminergic mechanisms of tic ontogeny. Systemic 5-HT_2A_R stimulation elevates cortical dopamine efflux, a process antagonized by volinanserin ^51,52^, implying that abnormal 5-HT_2A_R tone can bias dopaminergic signaling without requiring pathology to originate within striatal dopamine terminals. Such an arrangement accounts for dopaminergic signatures in TS while placing the initiating dysregulation upstream, at the level of prefrontal gain control.

Our postmortem findings indicate that, whereas 5-HT_2A_R protein levels did not differ between TS and control tissue in BA8 or BA9, BA10 exhibited an approximately threefold elevation in male TS samples. BA10 is a higher-order prefrontal subregion implicated in multitasking, prospective control, and the integration of internally generated goals with external demands ^53^. These functions map closely onto core clinical features of TS, including context-dependent fluctuations in tic suppressibility ^54,55^ and the worsening of symptoms during cognitive or emotional load ^56,57^. The male predominance of BA10 upregulation is also consistent with the well-established male bias in TS prevalence and, in many cohorts, symptom severity ^58–60^, raising the possibility that sex-dependent serotonergic regulation of prefrontal control contributes to the sexual dimorphism of TS. Notably, structural MRI studies have reported reductions in prefrontal gray matter that correlate with tic severity ^61^. When interpreted alongside the present molecular and pharmacological data, these observations suggest that elevated 5-HT_2A_R expression in BA10 may contribute to tic ontogeny by amplifying state-dependent fluctuations in cortical excitability, weakening top-down inhibitory control over motor output and other behavioral responses.

A key strength of the present work is the replication of anti-tic efficacy across two mechanistically distinct TS models. The D1CT-7 model, generated by inserting a cholera toxin gene under the D_1_ dopamine receptor promoter, exhibits cortical neuropotentiation and key motor features similar to tics ^36^. In this study, transitioning the line from a Balb/c to a mixed Balb/c–C57BL/6 background improved surgical viability but attenuated spontaneous tic expression, requiring mild stress (spatial confinement) to elicit robust jerks ^41^. Importantly, prior work demonstrated that tics in D1CT-7 mice are reduced by the 5-HT2 receptor antagonist ritanserin ^62^, a finding fully consistent with the present results. Additionally, both systemic and intra-mPFC administration of pimavanserin and volinanserin suppressed tic-like jerks, while intrastriatal administration was ineffective, demonstrating that local prefrontal 5-HT_2A_R blockade is sufficient to abolish tics, supporting a cortical locus of pathophysiological control. Complementary to these findings, the CIN-d model ^37^, which recapitulates partial loss of striatal cholinergic interneurons seen in TS postmortem studies ^63,64^, also displayed stress-induced tic-like behaviors that were normalized by the same 5-HT_2A_R antagonists. This convergence supports generalizability and suggests that prefrontal 5-HT_2A_R signaling may represent a final common pathway through which heterogeneous upstream perturbations influence tic expression.

Our pharmacological data further indicate a functional dissociation between simple and complex tic-like motor outputs in mice. Tic-like jerks were readily induced by 5-HT_2A_R stimulation and robustly suppressed by antagonism, supporting the conclusion that 5-HT_2A_R signaling is both necessary and sufficient for these brief motor events. In contrast, grooming stereotypies were reduced by 5-HT_2A_R antagonists but not reliably induced by low-dose agonist stimulation, suggesting that 5-HT_2A_R activity may be necessary, but not sufficient, for the expression of more complex motor sequences. This distinction is conceptually aligned with the clinical differentiation between simple tics, which consist of rapid, fragmentary motor outputs, and complex tics, which involve longer, patterned action sequences. Defining what constitutes a fully representative “tic-like” manifestation in rodents is inherently challenging ^65^, given species-specific motor repertoires. Within this framework, tic-like jerks may represent the most face-valid proxy for simple tics, whereas stereotyped grooming may reflect a broader category of repetitive or perseverative behaviors that partially overlap with, but are not identical to, complex tics.

Several mechanistic possibilities may explain why 5-HT_2A_R stimulation preferentially elicits jerks but fails to consistently trigger stereotypies at low doses. One possibility is that parvalbumin-positive interneurons in the striatum play a critical role in constraining 5-HT_2A_R-driven cortical excitation within a single motor loop, thereby favoring brief, focal motor fragments. Alternatively, higher levels or broader spatial recruitment of 5-HT_2A_R-dependent excitation may be required to overcome inhibitory gating and allow excitation to propagate into multiple action channels, resulting in more complex stereotyped behaviors. In this scenario, stereotypies would emerge only when excitatory drive exceeds a higher threshold or engages additional circuit elements. Future studies will be required to directly test these hypotheses and to determine whether the transition from simple jerks to complex stereotypies reflects quantitative increases in excitatory gain, qualitative changes in recruited cell types, or both.

Beyond tic suppression, a particularly important translational implication of 5-HT_2A_R-targeted therapy is its potential to address aggression and irritability, which affect a substantial proportion of TS patients and contribute disproportionately to impairment and family burden ^66,67^. Aggression in TS is strongly associated with higher tic severity and reduced social functioning ^68^. In the most severe presentations, often termed “malignant TS,” uncontrolled aggression, self-injury, and suicidality can necessitate emergency intervention and substantially increase morbidity ^60,69^. Current pharmacological strategies for TS-associated aggression rely on atypical antipsychotics such as risperidone and olanzapine ^20,21^, which combine D_2_ receptor antagonism with strong 5-HT_2A_R blockade ^70^. While these agents reduce tics and aggression, their utility is constrained by weight gain, metabolic disturbances, and sedation. These effects are largely attributable to the blockade of 5-HT_2C_ and histamine H_1_ receptors ^71–74^. These limitations are especially consequential in pediatric populations, where they compromise compliance ^75,76^. The present findings, showing that selective 5-HT_2A_R antagonism attenuates aggression in a TS-relevant model, support the inference that 5-HT_2A_R blockade itself contributes materially to controlling behavioral dysregulation. In line with this interpretation, previous studies have shown that elevated expression of 5-HT_2A_Rs in the PFC are associated with behavioral disinhibition and high impulsivity in rats ^77,78^. Furthermore, 5-HT_2A_R antagonists have been shown to reduce impulsivity in animal models ^79^.

The PFC, a key center for emotional control and social behavior, plays a central role in aggressive manifestations ^80,81,81^. Increased 5-HT_2A_R expression in the PFC of TS patients, as shown here, parallels findings in individuals with antisocial or aggressive traits ^82–84^ and supports a shared mechanism linking prefrontal serotonergic dysregulation to disinhibited behaviors. Notably, genetic polymorphisms in the *HTR2A* gene (encoding 5-HT_2A_R) have been associated with aggression and impulsivity ^85–87^. Additionally, 5-HT_2A_R antagonism elicits anti-aggressive effects across disorders characterized by irritability and behavioral disinhibition ^88^, plausibly through modulation of prefrontal-limbic circuitry regulating emotional reactivity, impulse control, and tic suppression. Accordingly, 5-HT_2A_R antagonists may provide dual benefit by stabilizing prefrontal regulation of both motor outputs and emotionally driven behavioral bursts. Aggressive behavior in TS has also been associated with coprophenomena ^89^, the involuntary expression of socially inappropriate words and gestures ^90^, which has a markedly negative impact on quality of life ^91^. It is therefore plausible that cortical 5-HT_2A_R overexpression contributes not only to elevated tic severity but also to these extreme disinhibitory symptoms. Future studies should determine whether prefrontal 5-HT_2A_R expression correlates with the presence and severity of coprophenomena, and whether selective antagonists such as pimavanserin are particularly effective in mitigating these disabling features.

In our experiments, pimavanserin did not impair general locomotor activity in D1CT-7. Consistent with these data, this drug led to very limited side effects in TS patients ^25^, and broader tolerability data report mild adverse events such as nausea or edema in fewer than 10% of patients ^26,92^. This profile compares favorably with existing TS pharmacotherapies. First-line treatments, such as antipsychotics and α2 adrenergic agonists effectively suppress tics but are limited by their significant adverse effects: for example, D_2_ receptor antagonists cause extrapyramidal symptoms, while atypical antipsychotics alter metabolism and α2 agonists cause sedation and hypotension ^76,93^ ^94^. In contrast, the highly selective mechanism of action of pimavanserin reduces the likelihood of extrapyramidal symptoms, sedation, and metabolic effects ^95^. Selective 5-HT_2A_R antagonism could therefore offer a mechanistically targeted alternative, particularly for patients intolerant of dopaminergic blockade or who display rage attacks.

Several limitations warrant acknowledgment. First, the postmortem dataset was predominantly male, reflecting both the male preponderance of TS and constraints inherent to brain bank availability. Given that TS may present differently in females, including later symptom onset and distinct comorbidity profiles ^96^, future studies are warranted to determine whether prefrontal 5-HT_2A_R upregulation generalizes across sexes. Second, although the two mouse models provide convergent evidence, neither fully recapitulates the genetic, developmental, and environmental complexity of human TS, nor captures phenomena such as complex premonitory urges. Future work should incorporate additional models, including those harboring TS-associated risk variants such as our *Celsr3* mutant ^97^, and should establish whether prefrontal 5-HT_2A_R upregulation precedes tic onset or emerges as a consequence of chronic symptomatology. Third, we exclusively examined the effects of acute drug administration. While this approach is valuable for establishing proof-of-concept efficacy, it does not address whether therapeutic benefits would be sustained under chronic treatment conditions. Prolonged 5-HT_2A_R antagonist exposure is known to induce compensatory receptor upregulation, whereas repeated agonist administration typically promotes receptor downregulation and desensitization ^98^. Such neuroadaptive processes could attenuate, potentiate, or qualitatively alter the behavioral effects observed acutely. Future studies employing chronic dosing paradigms will be essential to determine the durability of tic suppression and to identify any emergent tolerance or rebound phenomena. Fourth, the cellular substrates underlying 5-HT_2A_R signaling within medial PFC remain incompletely characterized. While our interpretation favors a primary implication of 5-HT_2A_Rs in the pyramidal cells, these receptors are also expressed on GABAergic interneurons, including parvalbumin-positive cells ^99,100^. Activation of these distinct populations may exert opposing effects on local circuit excitability and downstream output, yet our current data do not resolve which cell types actually show an overexpression of 5-HT_2A_Rs and drive the observed behavioral phenotypes. Elucidating these contributions will require cell-type-specific genetic manipulations, such as conditional receptor knockouts or chemogenetic approaches, combined with projection-resolved circuit mapping to determine how 5-HT_2A_R-expressing neurons in medial PFC influence cortico-striatal and cortico-limbic pathways implicated in tic generation. Finally, our behavioral assessments focused on tic-like movements and did not include measures of attentional function or response inhibition. Future studies incorporating tasks such as the five-choice serial reaction time task or stop-signal paradigms would clarify whether 5-HT_2A_R antagonism confers broader cognitive benefits beyond tic suppression, thereby strengthening the translational relevance of these findings.

These caveats notwithstanding, this study provides the first combined human–animal evidence linking tic pathogenesis to increased 5-HT_2A_R expression in the PFC and demonstrating that targeted antagonism suppresses both tics and aggression across species. These findings position cortical 5-HT_2A_Rs as promising therapeutic targets for TS and potentially for broader phenotypes characterized by behavioral disinhibition and impulsive aggression.

## ACKNOWLEDGMENTS

We thank Caterina Branca for editorial suggestions. This study was partially supported by NIH grants R21 NS125519 and R21 NS108722 (to M.B.). Postmortem human samples were obtained through the NIH NeuroBioBank from Harvard University Brain Tissue Repository.

## ETHICS STATEMENT

Tissues were obtained from the Brain Tissue Repository at Harvard University through the NIH NeuroBioBank program. All brain tissues were procured, stored, and distributed according to state and federal guidelines and regulations involving consent, protection of human subjects, and donor anonymity.

