## Supplemental Figure 1 for "Prefrontal 5-HT_2A_ receptors directly contribute to tic ontogeny: translational evidence"

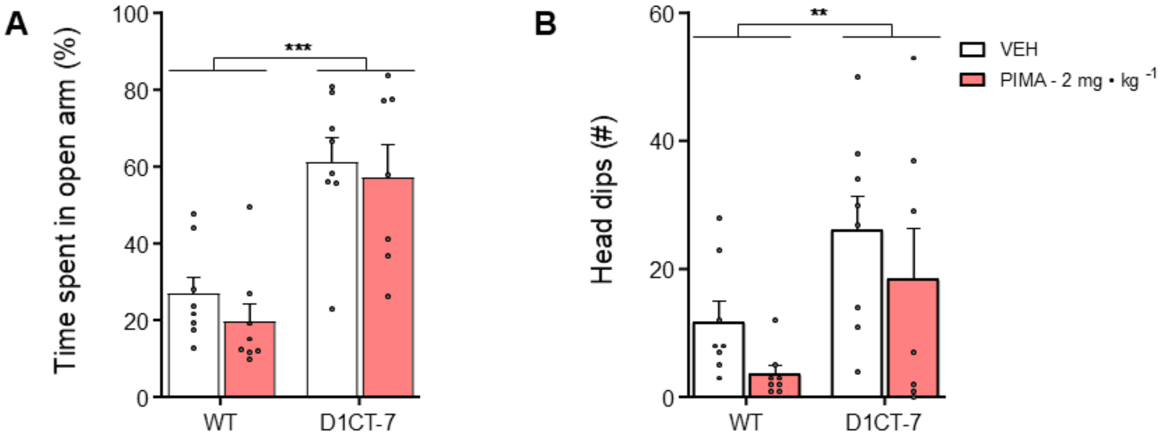


**Systemic administration of pimavanserin (PIMA; 2 mg·kg$^{-1}$) does not produce anxiolytic-like effects in the Elevated Plus Maze.** Two-way ANOVA revealed a significant main effect of genotype on open-arm time ($F(1,27) = 34.29$, $P < 0.0001$), with no significant effects of treatment ($F(1,27) = 0.85$, $P = 0.36$) or genotype × treatment interaction ($F(1,27) = 0.07$, $P = 0.80$). A comparable pattern was observed for head-dip frequency, with a significant effect of genotype ($F(1,27) = 8.79$, $P = 0.006$) and no effects of treatment ($F(1,27) = 2.57$, $P = 0.12$) or interaction ($F(1,27) = 0.003$, $P = 0.96$). All experiments were performed in male mice. Data are shown as mean ± SEM. ** $P < 0.01$; *** $P < 0.001$.
