## Supplemental Table 1 for "Prefrontal 5-HT_2A_ receptors directly contribute to tic ontogeny: translational evidence"

**SUPPLEMENTAL TABLE 1: Clinical and demographic characteristics of human postmortem brain tissue donors.** Postmortem tissue was obtained from 13 patients with Tourette syndrome diagnosis, as well as 13 controls matched for age and postmortem interval (PMI) from the NIH NeuroBioBank Brain Tissue Resource Center (Harvard University) through the Harvard Brain Tissue Repository.

| ID | SEX | AGE | PMI | Diagnosis |
|---|---|---|---|---|
| **1** | Male | 8 | 12.3 | Tourette's disorder |
| **2** | Male | 14 | 6.5 | Tourette's disorder |
| **3** | Male | 29 | 19 | Tourette's disorder |
| **4** | Male | 34 | 30.1 | Tourette's disorder |
| **5** | Male | 39 | 27 | Tourette's disorder |
| **6** | Male | 41 | 25.5 | Tourette's disorder |
| **7** | Male | 44 | 12.5 | Tourette's disorder |
| **8** | Male | 54 | 20.18 | Tourette's disorder |
| **9** | Male | 45 | 30.7 | Tourette's disorder, Myasthenia gravis, Obsessive-compulsive disorder |
| **10** | Male | 17 | 28.92 | No clinical diagnosis |
| **11** | Male | 24 | 13.1 | No clinical diagnosis |
| **12** | Male | 33 | 25.67 | No clinical diagnosis |
| **13** | Male | 35 | 25.67 | No clinical diagnosis |
| **14** | Male | 36 | 21.46 | No clinical diagnosis |
| **15** | Male | 43 | 14.68 | No clinical diagnosis |
| **16** | Male | 44 | 13.58 | No clinical diagnosis |
| **17** | Male | 47 | 26.83 | No clinical diagnosis |
| **18** | Male | 19 | 18.58 | No clinical diagnosis |
| **19** | Female | 20 | 35.75 | Tourette's disorder |
| **20** | Female | 48 | 14.8 | Tourette's disorder |
| **21** | Female | 55 | 13.42 | Tourette's disorder |
| **22** | Female | 59 | 25.5 | Tourette's disorder |
| **23** | Female | 42 | 20.34 | No clinical diagnosis |
| **24** | Female | 50 | 20.25 | No clinical diagnosis |
| **25** | Female | 55 | 21.66 | No clinical diagnosis |
| **26** | Female | 59 | 23.78 | No clinical diagnosis |
